# Cellar: Interactive single cell data annotation tool

**DOI:** 10.1101/2021.03.19.436162

**Authors:** Euxhen Hasanaj, Jingtao Wang, Arjun Sarathi, Jun Ding, Ziv Bar-Joseph

**Author notes:** Correspondence: Jun Ding, or Ziv Bar-Joseph.

## Abstract

Several recent technologies and platforms enable the profiling of various molecular signals at the single-cell level. A key question for all studies using such data is the assignment of cell types. To improve the ability to correctly assign cell types in single and multi-omics sequencing and imaging single-cell studies, we developed Cellar. This interactive software tool supports all steps in the analysis and assignment process. We demonstrate the advantages of Cellar by using it to annotate several HuBMAP datasets from multi-omics single-cell sequencing and spatial proteomics studies. Cellar is freely available and includes several annotated reference HuBMAP datasets.

**Availability:** https://data.test.hubmapconsortium.org/app/cellar

## Background

A number of large consortia are focused on profiling tissues, organs, and the entire human body at the single-cell level. Some examples include the Human BioMolecular Atlas Program (HuBMAP) [1], which aims to reconstruct a 3D map of the human body at the single-cell resolution, the Human Cell Atlas [2], The BRAIN initiative [3], the Human Tumor Atlas Network (HTAN) [4], and many more. These consortiums use several different technologies for studying the molecular composition of single cells. While the specific platforms vary, popular types of data collected by most consortia include single-cell RNA Sequencing, single-cell ATAC Sequencing [5], single-cell spatial transcriptomics [6], and single-cell spatial proteomics data [7]. In addition to the large consortia, several individual labs generate data using some or all of these modalities. One of the first questions researchers face when analyzing these single-cell datasets is cell-type assignment. Several different strategies and methods have been developed for the assignment of the various data types being profiled [8, 9, 10, 11, 12, 13]. While the methods differ in details, the most popular ones follow a standard set of steps. First, the dimension of the data is reduced. Next, the data is clustered to obtain groupings of similar cells. Then the clusters are displayed (usually by further reducing the dimension of the data). Finally, marker or enrichment analysis is performed to assign a cell type to each cluster. Each of these steps has been the subject of several different studies, each proposing a different analysis method ranging from optimal ways to reduce the dimension of different types of omics single-cell data [14] to methods for clustering such data [15] to supervised and unsupervised methods for determining cell types based on the set of genes expressed in each of the clusters [8].

To date, different groups from the same consortia and even the same group when processing multiple types of single-cell omics data use a different set of tools. This makes it hard to integrate and compare data from these groups, even for the same tissue and modality. It is even harder to integrate data from different modalities since researchers often use various assignment techniques, markers, and even cell-type naming conventions for analyzing their datasets. To enable large scale collaborations, integration, and comparisons across many different single-cell omics platforms and modalities, we developed a new, interactive and graphical cell-type assignment server termed Cellar. Cellar provides all the functionality needed for cell-type assignment and supports all the data types described above. As part of Cellar, we implemented many popular dimensionality reduction and clustering methods and several different visualization and embedding modules. Cellar offers the ability to transfer labels from previously annotated datasets to new data, use several established or user-defined ontologies for markers, or constrain the set of cell types assigned (for example, using an agreed list). Finally, we implemented components to support interactive and semi-automated assignments, including methods for constrained clustering, for joint analysis of scATAC-seq and scRNA-seq data, for visualizing markers and combination of markers, and more. Figure 1 shows a high-level graphical representation of Cellar’s workflow. We tested Cellar by annotating several different organs, datasets, platforms, and modalities from HuBMAP. These include 10x, SNARE-seq, scATAC-seq, and Codex spatial proteomics data. As we show, using Cellar, researchers were able to successfully assign cell types for all these datasets and combine cell-type assignment across modalities.

**Figure 1:**
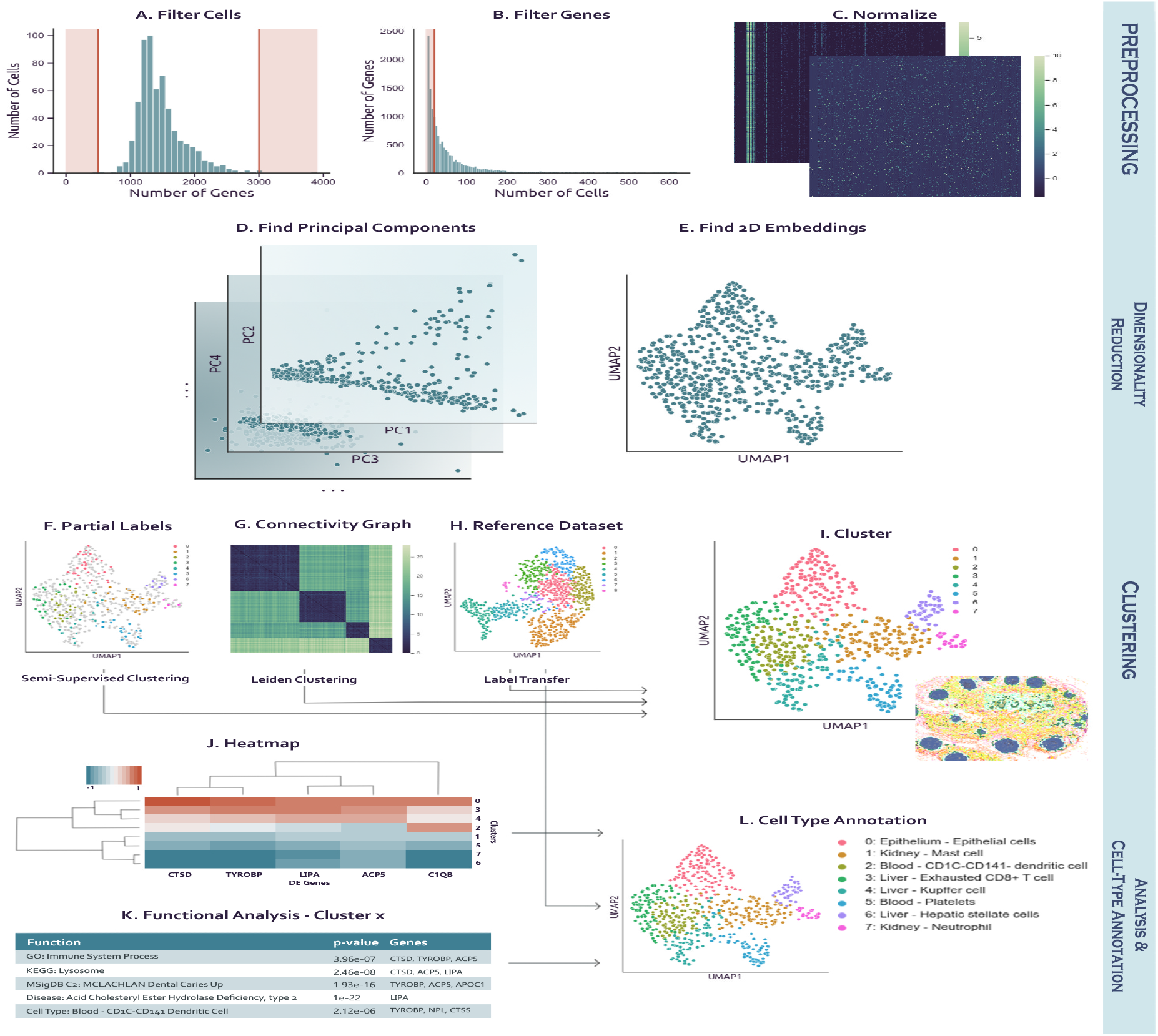
Cellar’s workflow. a, b, c: Preprocessing (optional). Cellar can filter cells based on the number of expressed genes and genes which are rarely expressed. Next the input is normalized. **d, e:** Dimensionality reduction and visualization. Several methods for dimensionality reduction are implemented as part of Cellar. The reduced data is then visualized by running another (possibly the same) dimensionality reduction method. **f, g, h, i:** Clustering. Cellar supports several unsupervised and semi-supervised clustering methods. It only implements supervised alignment methods. **j, k, l:** Cell type assignment. Cellar enables the use of several functional annotation databases for the assignment of cell types.

## Results

We developed Cellar, a web-server for interactive cell-type assignment in single-cell omics studies. Cellar supports a wide range of methods for all the steps involved in the process of cell-type assignment. As cell-type assignment often requires user input at various steps of the pipeline, Cellar adopts a semi-automatic solution where users have the opportunity to interact with the tool at each critical step (e.g., clustering, labeling). In the clustering step, the users have the opportunity to merge or split clusters based on their prior knowledge and expertise. In the labeling step, Cellar displays comprehensive information that can guide the annotation step (e.g., top Differential genes, enriched GO terms, enriched pathways, enriched Gene Set Enrichment Analysis (GSEA) functional terms, comparison with known cell-type markers, enriched diseases). All the clustering and labeling results can be saved and exported as an AnnData object [16]. This session file can be shared with others or loaded back into Cellar for further analysis.

### Cell-type assignment for scRNA-seq data

We used Cellar to annotate 15 scRNA-seq datasets (10x genomics) with an average of 7,500 cells from five different tissues (Kidney, Heart, Spleen, Thymus, Lymph node) [17], all of which can be found on the Cellar website. We demonstrate the functionalities of Cellar by discussing one of these datasets sampled from a spleen tissue (HBMP3 CC2) with 5,273 cells. First, Cellar performs standard preprocessing to filter out ‘bad-quality’ cells based on user-defined criteria (Methods). We then apply dimensionality reduction to the preprocessed and normalized expression matrix, followed by clustering and visualization of the resulting clusters. Initially, we obtained 18 clusters in total for this spleen dataset (Figure 2a). For each of those clusters, cellar identifies top differential genes (Figure 2d).

**Figure 2:**
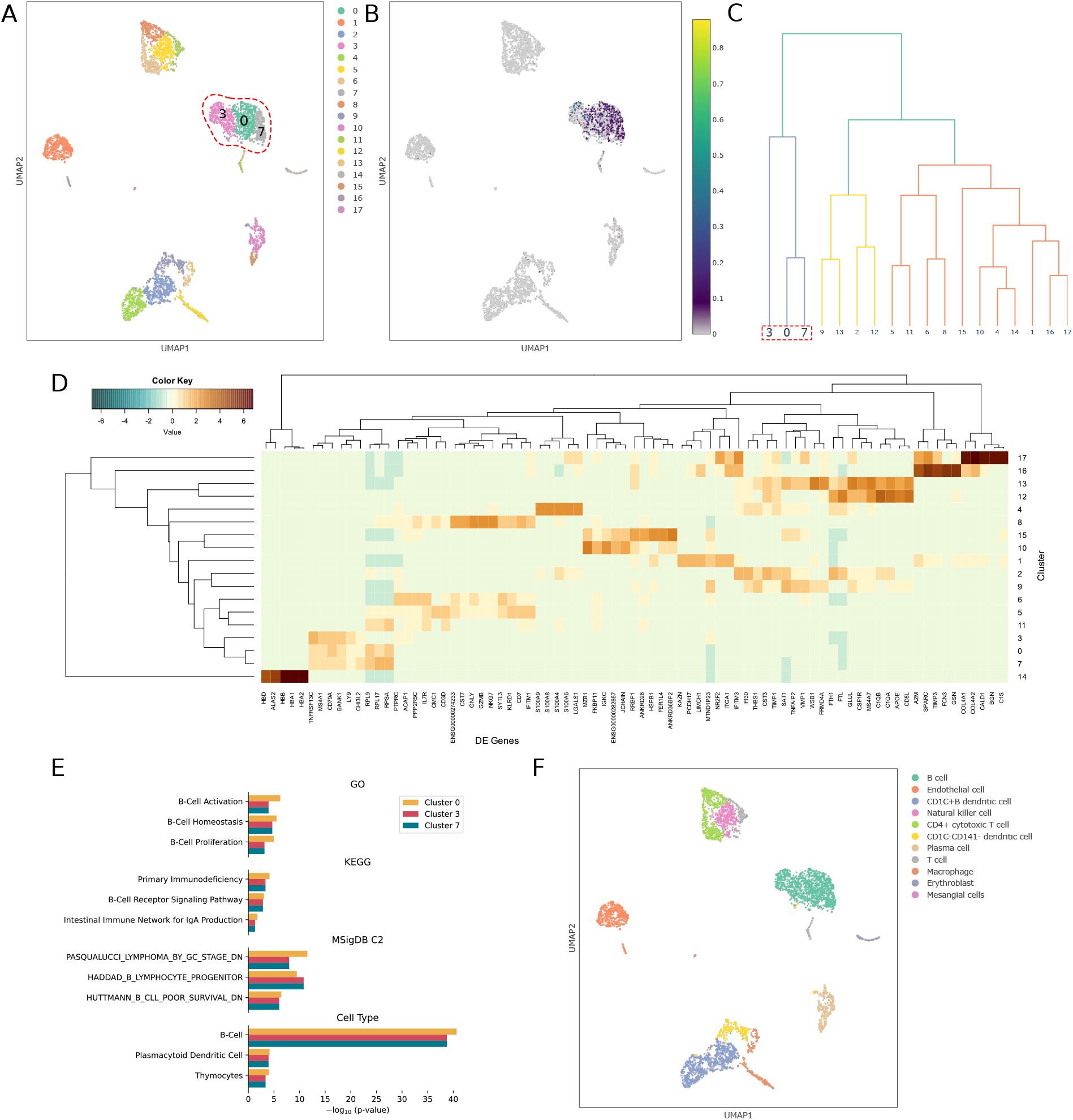
scRNA-seq example. **a**. Screenshot of a spleen single-cell dataset clustered using Leiden. Based on functional enrichment analysis, Cellar shows strong evidence that clusters 0, 3, and 7 (circled) consist of B-cells. **b**. A gene co-expression view of CD79A and TNFRSF13C. Both genes are known markers for B-cells and are highly co-expressed in the circled clusters. **c**. A dendrogram displaying the hierarchical relationship between the clusters, using the top 100 DE genes for each cluster. There is a strong relationship between clusters 0, 3, and 7. **d**. Top 5 DE genes for each cluster. **e**. Top functional analysis (GO, KEGG, MSigDB, Cell Type) entries for clusters 0, 3, and 7 and their corresponding p-value. There is strong support for the B-cell type. **f**. Example annotations for the spleen dataset obtained using Cellar. Based on the results of functional analysis, we merged several pairs of clusters, including 0, 3, and 7. The cell type was determined after considering p-values and consistency across different ontologies.

The number of top differential genes varies from 100 to 1000, with most clusters displaying hundreds. Users can apply functional enrichment analysis on those DE genes (GO term, Pathway, MSigDB, disease) to determine the potential functions and cell types associated with each cluster. For example, for clusters 0, 3, and 7, we show the top results for GO, KEGG, MSigDB, and Cell type functional analysis in Figure 2e. These clusters are enriched with GO terms “B-Cell Activation” and “B-Cell Homeostasis”, KEGG Pathway “B-Cell Receptor Signaling”, and MSigDB pathway “HADDAD B Lymphocyte Progenitor”, all of which suggest that clusters 0, 3, 7 are likely B-cells. This is further confirmed by looking at the cell type analysis where “B-Cell” is the top entry. Users can either merge these clusters or refine the specific annotation for each.

Cellar also supports the visualization of specific combinations of markers. Often multiple cell type makers are required to pinpoint the cell type accurately. For example, CD3+ and CD8+ are needed to find cytotoxic T-cells. To allow users to visualize cells that highly express multiple marker genes, we provide a gene co-expression view (Methods). Visualizing two marker genes for B-cells (Figure 2b) indicates that they are highly expressed in clusters 0, 3, and 7. Cellar enables users to manually or semi-automatically combine clusters, as shown in Figure 2f. Following a manual combination of clusters 0, 3, and 7, we observe that the top three DE genes for the merged cluster are CD79A, TNFRSF13C, MS4A1, which are all known markers for B-cells [18]. A complete list of cell types identified using Cellar is shown in Figure 2f.

### Cell-type assignment for scRNA-seq data using Label transfer

Cellar can transfer the cell-type annotations from already labeled data to new data from the same tissue. We used an annotated spleen dataset (HBMP3 CC3) to transfer labels to HBMP3 CC2 using SingleR [19]. The label transfer results are shown in Figure 3a. While most cells are assigned accordingly, there are a few low-quality annotations. Furthermore, several clusters’ functional analysis does not correspond to the assigned cell type (e.g., Hematopoietic stem cell, Plasmablast, Splenic fibroblast). The users can refine the annotation for these clusters or use the semi-supervised clustering methods provided in Cellar.

**Figure 3:**
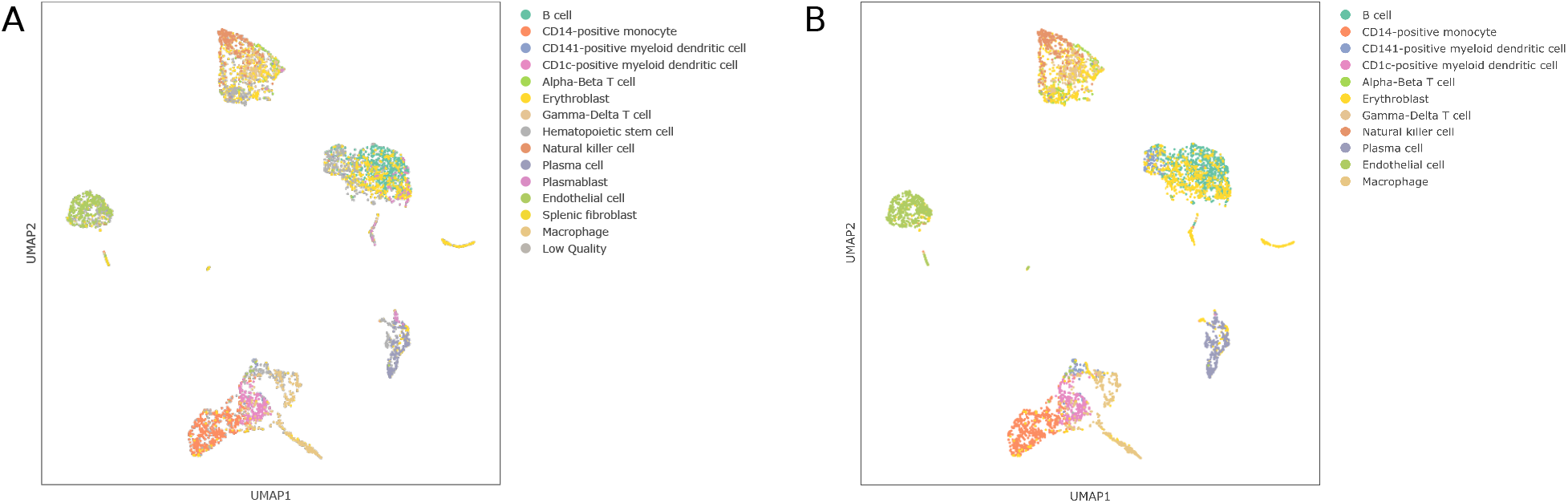
Label transfer example. **a**. The result of transferring labels from an annotated dataset of the same tissue to our HBMP3 CC2 spleen dataset using the pruned labels from SingleR. Low quality cells were not assigned a label. **b**. Revised assignments after using the semi-supervised clustering option. The cells belonging to the clusters: “Low Quality,” “Hematopoietic stem cell,” “Plasmablast,” and “Splenic Fibroblast” have been redistributed among other clusters using the Constrained k-Means algorithm. This increases the silhouette score from −0.03 to 0.1.

We used Cellar’s semi-supervised clustering tool. We specified that all the clusters but the low-quality ones should be preserved and decided to iteratively re-distribute any low-quality points to the nearest centroids via the Constrained k-Means algorithm (Methods). The results of semi-supervised clustering are shown in Figure 3b. The cluster assignment following label transfer achieves a silhouette score of -0.03, which is increased to 0.1 after running constrained clustering. Optionally, users may specify a number of clusters greater than the number of those preserved. These “new” clusters can then be annotated in a similar manner as we describe in the previous section.

### Cell-type assignment for scATAC-seq data

scATAC-seq [5] measures the genome-wide chromatin accessibility at the single-cell level, which profiles the epigenetic landscape for gene regulation that may be critical for determining cell types. Cellar handles scATAC-seq data in two different ways: cell-by-gene and cell-by-cisTopic. The former is based on the open chromatin accessibility associated with the nearby region of all genes ([-5KB, +1KB]). The latter relies on cisTopic [13] which uses Latent Dirichlet Allocation [20] to model cis-regulatory topics. The resulting cell-by-gene or cell-by-topic matrix is used for downstream analysis such as visualization and clustering.

We show an example of the former method on a public 10x scATAC-seq dataset consisting of Peripheral Blood Mononuclear Cells (PBMC) [21]. We obtain a cell-by-peak matrix that contains 9,277 cells and 80,234 peaks. For every protein-coding gene as specified in GENCODE v35 [22], we sum all the peaks whose range intersects the gene location and up to 5KB upstream and 1KB downstream. The cluster results are shown in Figure 4a.

**Figure 4:**
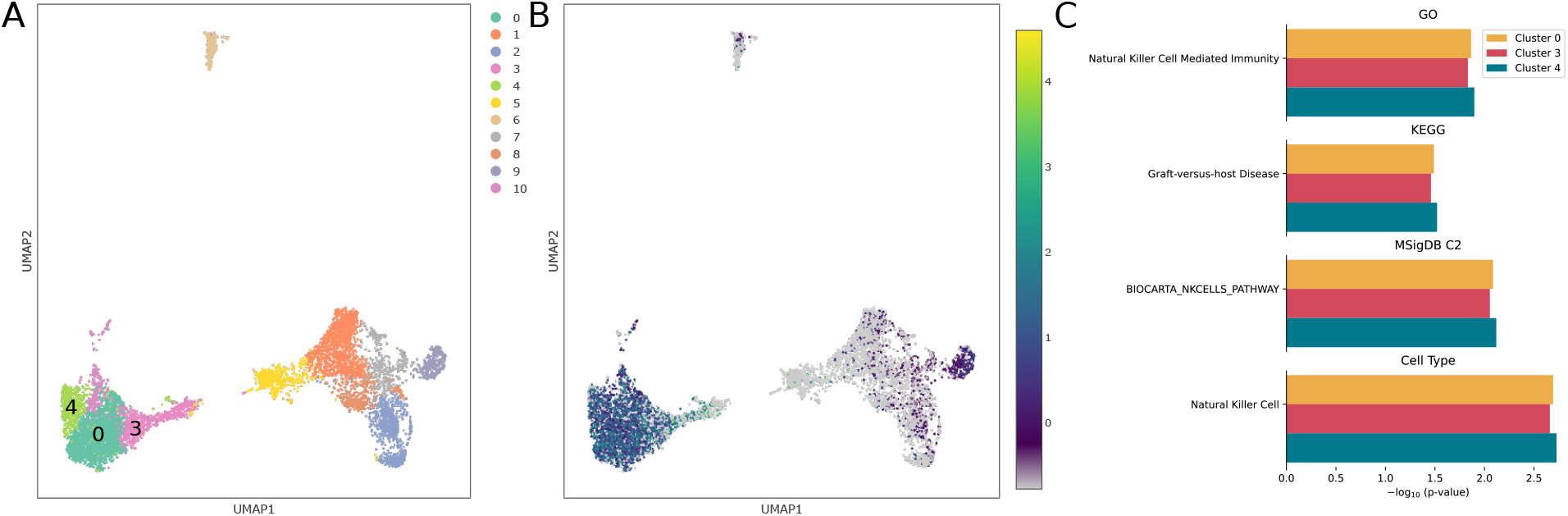
scATAC-seq example. **a**. Leiden clustering of 10k Peripheral Blood Mononuclear Cells scATAC-seq data. **b**. Activity levels for the KLRD1 marker gene present in clusters 0, 3, and 4. **c**. Top enriched functional terms for clusters 0, 3, and 4 and the corresponding p-value. The analysis is suggestive of the Natural Killer cell type.

We analyze clusters 0, 3, and 4. We run a t-Test to determine differentially expressed genes and use these to determine the cell type for these clusters. A list of functional terms and potential cell types given known markers is shown in Figure 4c. The KLRD1 gene, which is a known marker for natural killer (NK) cells [23, 24], is highly expressed in all these clusters (Figure 4b). Furthermore, functional enrichment analysis on these clusters provides strong support for the natural killer cell type (GO term “Natural Killer Cell Mediated Immunity”, KEGG pathway “Graft-versus-host Disease” [25], and MSigDB pathway “BIOCARTA NKCELLS PATHWAY”).

### Cell type assignment for single-cell spatial proteomics data

CO-Detection by indEXing (CODEX) is a spatial proteomics technology that provides information on protein levels at the single-cell resolution [26]. Typically, a CODEX dataset may consist of tens of thousands of cells, making data analysis very challenging. Here, we show that Cellar is capable of handling large CODEX datasets while providing convenient tools for analyzing both the expression profiles of cells and, at the same time, visualizing the spatial information. We use a lymph node dataset that contains 46,840 cells. After preprocessing, we reduced the dimensionality via UMAP to 10 dimensions, followed by Leiden clustering. The clustering results are shown in Figure 5a. Our experiments showed that PCA was not appropriate for analyzing CODEX data, while UMAP resulted in better separation of clusters. The corresponding tile for these cells with the projected cluster annotations is presented in Figure 5b. Given the small number of proteins profiled in this dataset (18), not all clusters could be assigned to unique types, though several have been assigned based on DE gene analysis in Cellar. Cellar matches the cell colors in the clustering and spatial images, making it easier to identify specific organizational principles and their relationship to the profiled cell types. As can be seen, the B-Cell clusters are surrounded by T-cells and other cells types in the lymph. The B-Cell clusters also contain a subset of proliferating cells.

**Figure 5:**
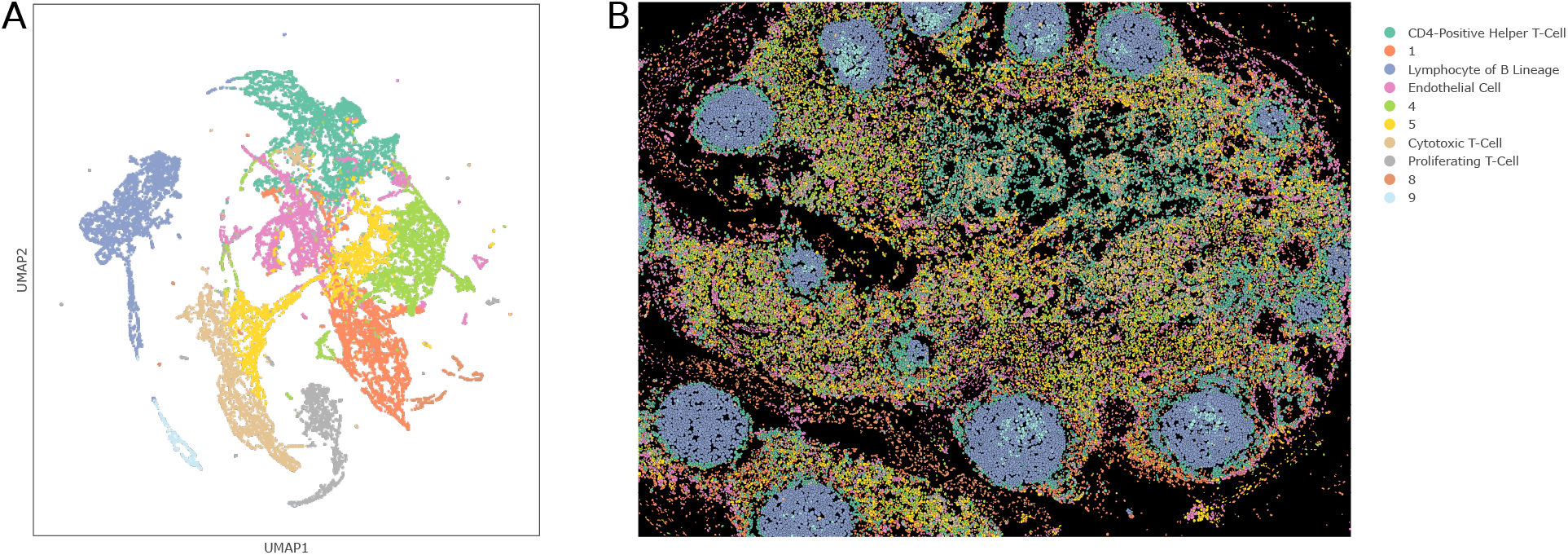
CODEX example. **a**. Visual representation of a lymph node CODEX dataset. Data was first reduced by UMAP and clustered by Leiden. **b**. Projection of the assignments on the spatial codex image that can be visualized side-by-side in Cellar. The colors correspond to the colors of the clusters in **a**. Not all clusters could be assigned to unique cell types given that only a few ten protein levels are measured, though several have been assigned based on the differential gene analysis in Cellar. The B-Cell clusters are surrounded by T-cells and other cells types in the lymph. The B-Cell clusters also contain a subset of proliferating cells

### Cell type assignment combining different data types

We have described how to analyze several different data types. Cellar offers the possibility to analyze multiple datasets simultaneously, not necessarily of the same type. Recent technologies allow studies that profile multiple types of biomolecules at the single-cell level. Cellar can annotate such data while taking into account the relationship between the values obtained for each modality. To demonstrate this, consider a SNARE-seq dataset [27] where both the transcriptome and chromatin accessibility of 32,278 cells is profiled. We first run cisTopic on the chromatin data and determine cluster assignments by running Leiden on the inferred cis-regulatory topics (Figure 6a). We use these labels to visualize the expression data in Figure 6b. This can be easily achieved using Cellar’s side-by-side mode, which allows both datasets to be visualized and analyzed simultaneously.

**Figure 6:**
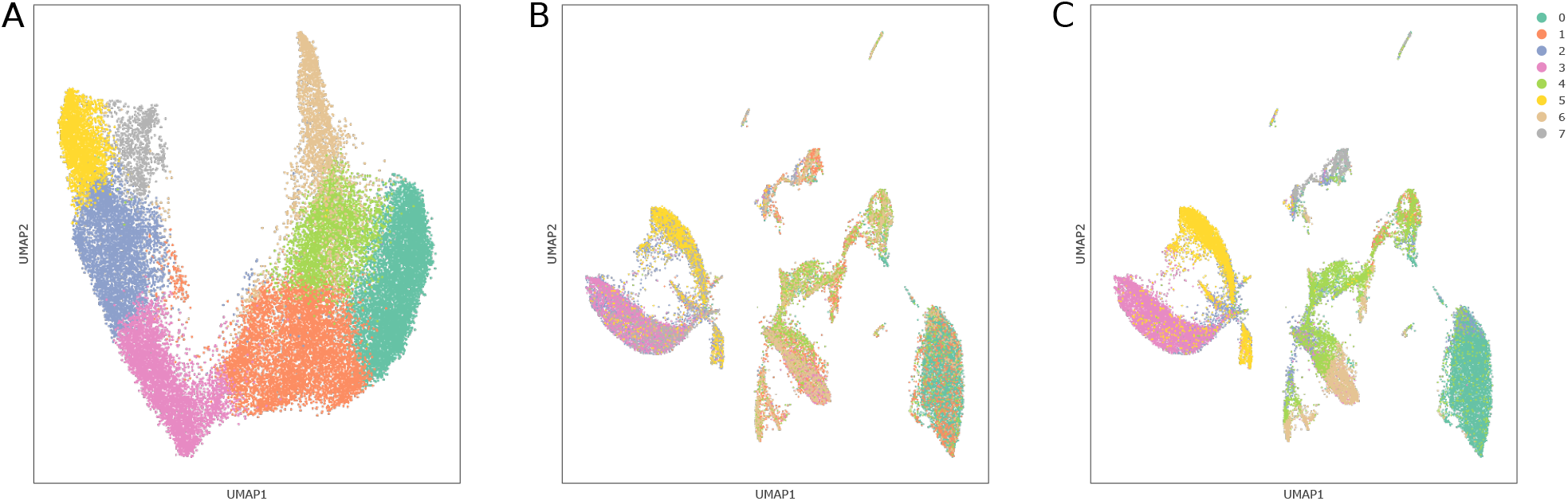
SNARE-seq example. Joint chromatin accessibility and expression data analysis in Cellar. **a**. Leiden clustering on the cell-by-cistopic matrix obtained by running cisTopic on the chromatin data. **b**. Visual representation of the corresponding expression data. Cell colors match those in **a. c**. Cluster assignment after a run of Constrained k-Means with 8 clusters. The resulting cluster configuration serves as a basic example to demonstrate how semi-supervised clustering can improve cluster homogeneity.

Using these labels, we decide to test the effect of semi-supervised clustering on the expression data. We run Constrained k-Means with 8 clusters and choose to preserve clusters 0, 3, 4, 5, which have better homogeneity (Figure 6c). This increases the silhouette score from -0.045 to 0.035. Cellar identifies differential genes, and we apply functional enrichment analysis to determine cell types for each of the clusters. For example, cluster 0 is enriched with GO term “Apical Plasma Membrane” (p-value=1e-4) which signifies the presence of Proximal Tubule Cells [28]. The cell type returned by cellar is also Proximal Tubule Cells (p-value=2.48e-6).

## Discussion

Cell type assignment has emerged as a major step in all single-cell based studies. While the vast majority of single-cell data is currently obtained from scRNA-seq studies, newer technologies are now generating several other types of single-cell data. These include epigenomics single-cell assays (for example, scATAC-seq), spatial transcriptomics, and spatial proteomics).

While several methods have been developed to aid researchers in the assignment of cell types for single-cell data, to date, these were focused on a single method (clustering, alignment, etc.) or modality (scRNA-seq, spatial transcriptomics, etc.). Recent studies require the use of multiple such methods and, in some cases, profile multiple types of scRNA-seq data. Cellar addresses these issues by providing a one-stop interactive platform for the analysis, visualization, and assignment of cell types in single-cell studies.

Cellar has already been used to annotate several different types of single-cell data from multiple platforms and tissues. In addition to serving as a proof of concept for the use of Cellar, our annotated datasets (mostly from HuBMAP) serve as a reference for transferring labels to other datasets using our implemented alignment methods. For tissues not currently supported by our HuBMAP annotated datasets, Cellar provides several external functional enrichment datasets that, combined with the user’s knowledge about specific markers, will hopefully make such assignment easier.

## Conclusions

Cellar is an easy to use, interactive, and comprehensive software tool for the assignment of cell types in single-cell studies. Cellar supports several types of molecular sequencing and imaging data and implements the most popular methods for visualization, clustering, and analysis. If fully integrated with the most popular functional annotation databases, we hope that Cellar would improve the accuracy and ease of cell-type assignment in single-cell studies. Cellar and the currently annotated data are freely available at [29].

## Methods

### Data upload and preprocessing

Cellar supports multiple scRNA-seq formats (either as raw read counts or as a normalized expression matrix in cell-by-gene format). Users can upload an expression matrix in h5ad format [16] which allows high compression levels. Cellar also accepts direct outputs from the 10x Genomics CellRanger pipeline as a gzip file, or even simple CSV/TSV (comma/tab separated) expression files. After loading the expression matrix into Cellar, users can choose whether to preprocess the data or not. Cellar provides multiple options for filtering cells and genes with bad quality. These include a customized cutoff on the number of expressed genes in each cell, mitochondrial read percentage, and the number of cells in which a gene is expressed. Users can also filter genes based on the dispersion and max expression value. Following these steps, the gene expression matrix is normalized and log1p transformed.

### Dimensionality reduction

Following preprocessing, the number of genes usually remains high, with a significant amount expressed at low levels in most cells. For this reason, clustering is typically preceded by a dimensionality reduction step which often improves the accuracy of clustering algorithms and makes the computations more tractable. Depending on the transformation map, such techniques fall into two categories, linear and non-linear. Cellar supports several options from both categories, including Principal Component Analysis (PCA) [30], Diffusion Maps [31], and Uniform Manifold Approximation and Projection (UMAP) [32], among others. PCA is a linear method that projects the data to a set of principal components (PC) that preserve as much of the variance as possible. In Cellar, the number of PCs can be specified by the user, or it can be automatically inferred by employing the elbow heuristic. For the latter, we apply the kneedle algorithm [33] to the graph of explained variance to determine the “elbow” of the curve. Diffusion Maps is a non-linear method that obtains the geometric structure of the data by focusing on local distances between points viewed at different scales. [34] argue in favor of using Diffusion Maps for analyzing single-cell data, as they are robust to noise and their nature is well-suited to the non-linear cell differentiation process. UMAP is another non-linear dimensionality reduction technique that tries to preserve the global structure of the data by finding a lower-dimensional approximation of the manifold that the data lies in. Other methods provided by Cellar include Multidimensional Scaling (MDS) [35], Isomap [36], and Truncated SVD [37] for efficiently working with sparse matrices [38].

In addition to improving the clustering of scRNA-seq data, dimensionality reduction is also useful for visualization and display. By projecting the high dimensional (often thousands of dimensions) datasets into a two-dimensional space, those complex datasets can be visually read by a human, which can help better understand the abstract high-dimensional space. All the above methods can be used for such visualization purposes.

### Clustering

Clustering is a crucial step when analyzing single-cell data because it uncovers groupings of similar cells which are then used for downstream analysis, such as identifying cell types. Cellar supports both unsupervised and semi-supervised clustering algorithms. On the unsupervised side, the default clustering method is Leiden [39], a community detection algorithm that uses an iterative procedure to merge small communities into larger ones such that the local modularity is increased. Leiden is also a popular choice in other packages [16, 40, 41]. Leiden takes as input a connectivity graph which we construct by using a nearest neighbors approach. The cluster configuration generated by Leiden is very sensitive to its resolution hyperparameter which is why we allow the users to tune its value. Our experiments show that Leiden generally overestimates the number of clusters when resolution equals 1. However, we found this to be a useful property as the users can use their domain knowledge to merge some of these clusters or rely on semi-supervised learning algorithms to generate better cluster configurations.

Cellar also provides other classical unsupervised learning algorithms such as k-Means, k-Medoids [42], Spectral Clustering [43], and Agglomerative Clustering. These clustering algorithms require the user to specify the number of clusters, *k*, which is typically not known in advance. To overcome this limitation, Cellar can run multiple clustering instances using different *k* and select the optimal number of clusters that achieves the highest Silhouette Score [44]. Cellar also provides an Ensemble Clustering method based on Hypergraph Partitioning [45, 46], which allows the users to explore various ensembles of clustering methods.

In addition to the unsupervised clustering methods mentioned above, we included two semi-supervised extensions of k-Means first presented in [47]. Cellar offers users the opportunity to manually define clusters by interacting with the plot and re-configuring the cluster assignments via selection tools. Often these manually defined clusters fail to satisfy optimality conditions, such as the k-Means objective. To address this issue, one can use these manually defined clusters as seeds for a new run of k-Means (Seeded k-Means), or specify a set of clusters they wish to “preserve” while (re-)clustering the remaining ones (Constrained k-Means, Methods). These semi-supervised clustering algorithms also improve label transfer as discussed below. Formally, given cells *c*_*i*_ ∈ ℝ^*d*^ and existing labels *y*_*i*_ ∈ [*k*], where *i* ∈ [*N*], given k-Means centroids *g*_*j*_ ∈ ℝ^*d*^ for *j* ∈ [*k*], and a set 𝒫 ⊂ [*N*] of cell IDs that the user wishes to preserve, we modify the assignment step (E-step) of k-Means as follows:

**for** *i* = 1, …, *N* **do**

**if** *i* ∉ 𝒫 **then**

Set *y*_*i*_ = arg min_*j*_ *d*(*c*_*i*_, *g*_*j*_)

where *d*(·, ·) is the Euclidean distance. The M-step does not change. We repeat both steps until convergence.

### Cell type assignment using markers

To determine cell types, Cellar first identifies a list of differentially expressed (DE) genes (potential marker genes) that are uniquely expressed in a selected subset of cells (any cluster or user-defined group of cells). Statistical tests such as the Wald test [48], the Welch’s t-test [49], and the Wilcoxon rank-sum test [50] are implemented within Cellar to identify differential genes (markers). Once DE genes are identified, Cellar allows users to perform a list of functional enrichment analyses on the obtained DE genes to infer the potential cell type and functions for the selected subset. The list of functional analyses includes GO [51], KEGG [52], MSigDB [51], Disease [51]. We also curated a list of known cell-type marker genes from [53, 54]. Using a hypergeometric test on the DE genes and the known cell type markers, Cellar can provide a list of potential cell type predictions for the selected subset. As the curated cell type list is far from complete, Cellar also allows users to upload a list of user-defined cell type markers. Besides the aforementioned cell type marker enrichment based cell type identification, cellar also enables the users to infer the cell types based on the expression of a few well-defined cell type markers. For example, T cells can be marked by CD3. By exploring the expression level of those well-defined marker genes in all clusters, the users could potentially determine the cell types. However, many cell types could not be marked by a single marker gene. Often a combination of cell type markers are needed to uniquely distinguish one cell type from another. Here we also provided such a co-expression based cell coloring scheme in cellar. Gene co-expression is used to visualize the co-expression levels of two genes at the same time. Formally, if we let *m*_*i*_ and *M*_*i*_ denote the lowest and highest expression values for gene *i* where *i* ∈ {1, 2}, we color each cell *c* as

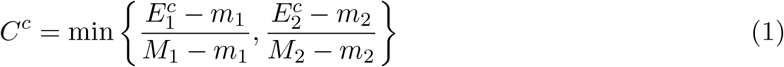

where *C*^*c*^ is the color of cell *c* and 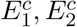 are the expression levels of gene 1 and 2 for cell *c*, respectively. We first map the ranges to [0, 1] to allow a fair comparison between the two genes, and we choose the minimum between the two, because the minimum satisfies the nice property min{0, *x*} = min{*x*, 0} = min{0, 0} = 0 for positive *x*. This is a desirable property since in all three cases we would want the “co”-expression levels to equal zero.

### Cell type assignment using label transfer

Label transfer or alignment utilizes previously annotated datasets (reference) to annotate new data. Alignment is done by finding similarities between cells in the reference and target datasets. Several methods have recently been developed for this task, and Cellar currently supports Scanpy Ingest [16] and SingleR [19]. Cellar includes several reference datasets based on HuBMAP annotated scRNA-seq, scATAC-seq, and spatial proteomics data. A side-by-side mode has been implemented to ease the comparison between the reference and target data as well as perform exact label transfer when the two datasets share cell IDs (e.g., in the case of SNARE-seq [27]).

## Declarations

### Ethics approval and consent to participate

No ethical approval was needed for this study.

## Consent for publication

Not applicable

## Availability of data and materials

The cellar software is publicly available on GitHub at [55]. The users can build a standalone version cellar following the instructions provided in the above GitHub page. The public cellar web-server is freely available at [29]. The cellar usage details can be found in the user guide at [56]. The datasets hosted on the cellar server are all from HuBMAP initiative [17].

## Competing interests

The authors declare that they have no competing interests

## Funding

Work partially funded by the National Institutes of Health (NIH) grants 1R01GM122096 and OT2OD026682 to Z.B.J..

## Authors’ contributions

ZBJ and JD designed the software. EH developed and implemented the back-end dimensionality reduction, clustering, and cell type annotation methods under the supervision of ZBJ and JD. EH, JW, and AS contributed to the implementation of the front-end interactive visualizations. JW and AS also contributed to the implementation of enrichment analysis of identified signature genes. All authors contributed with manuscript writing. The authors read and approved the final manuscript.

## Acknowledgements

Not applicable

## Notes

https://data.test.hubmapconsortium.org/app/cellar

### Competing Interest Statement

The authors have declared no competing interest.

